# Receptor levels determine binding affinity of WNT-3A to Frizzled 7 in a colorectal cancer model

**DOI:** 10.1101/2022.07.04.498383

**Authors:** Lukas Grätz, Joanna J. Sajkowska-Kozielewicz, Janine Wesslowski, Katja Petzold, Gary Davidson, Gunnar Schulte, Paweł Kozielewicz

## Abstract

WNT binding to Frizzleds (FZD) is a crucial step that leads to the initiation of signalling cascades governing multiple processes during embryonic development, stem cell regulation and adult tissue homeostasis. Recent efforts have enabled us to shed light on WNT-FZD pharmacology in overexpressed HEK293 cell systems. However, it is important to assess ligand binding at endogenous receptor levels as there might be differential binding behaviour in a native environment. Here, we focus on one FZD paralogue: FZD_7_, and study its interactions with WNT-3A in a CRISPR-Cas9-edited SW480 colorectal cancer model. SW480 cells were CRISPR-Cas9-edited to insert a HiBiT-tag on the N-terminus of FZD_7_, preserving the native signal peptide. Subsequently, these cells were used to study eGFP-WNT-3A association to endogenous and overexpressed HiBiT-FZD_7_ using NanoBiT/BRET to measure ligand binding and quantification of NanoBiT-emitted luminescence to assess receptor internalization. eGFP-WNT-3A bound to endogenous HiBiT-FZD_7_ with significantly higher *k*_on_ and with lower *K*_d_ than to overexpressed receptors. Importantly, as the fluorescent probe is an agonist, experiments performed in cell lysates demonstrated that eGFP-WNT-3A/HiBiT-FZD_7_ binding assessment is not altered by receptor internalization. In conclusion, binding affinities of eGFP-WNT-3A to HiBiT-FZD_7_ decreased with increasing receptor concentrations suggesting that HiBiT-FZD_7_ overexpression fails to recapitulate ligand binding behaviour in a (patho-)physiologically relevant context where endogenous receptor expression levels are lower.

## 1. INTRODUCTION

The ten Frizzleds (FZD_1-10_) and Smoothened (SMO) are G protein-coupled receptors (GPCRs), which form the class F of the GPCR superfamily [1, 2]. While endogenous sterols bind SMO, the 19 different Wingless/Int-1 (WNT) lipoglycoproteins are the main macromolecular ligands of FZDs [2, 3]. WNT-FZD interactions, which are the focus of this work, occur between the protein ligand and the extracellular cysteine-rich domain (CRD) of the receptor [4–7]. In addition to FZD binding, WNTs also associate with other WNT receptors, e.g. from the lipoprotein-receptor-related-protein family (LRPs) [8]. The binding of WNTs to their receptors initiates β-catenin-dependent or β-catenin-independent signalling, which regulate multiple processes during embryonic development, stem cell regulation, and adult tissue homeostasis [9, 10]. Furthermore, aberrant WNT signalling is associated with tumorigenesis and other pathologies [11, 12]. Along these lines, the last 30 years of research have provided substantial knowledge about WNT-induced signalling in healthy and pathophysiological conditions, but we have only recently started to understand the mechanistic and pharmacological details of WNT-FZD interactions [9, 11, 13–17]. To this end, FRET- and BRET-based approaches applied to live HEK293 cells allowed analysis of WNT-CRD interaction dynamics [18], WNT-induced conformational changes in FZDs [19–21], and to quantifiably define ligand-receptor binding using full-length receptors [13, 14]. These studies, however, were based on overexpressed recombinant proteins – an approach that may fail to accurately simulate physiologically relevant conditions [22, 23].

To overcome these limitations, endogenous proteins can be tagged using genome modification with CRISPR-Cas9 [24]. This technology has been employed with success in a number of studies designed to understand the biology of GPCRs [25] as well as a study on the co-receptor for Wnt/β-catenin signalling, LRP6 [26]. Among these, several studies have reported on NanoBRET ligand binding to endogenous class A GPCRs, mostly in HEK293 cells [27–32]. Here, as a continuation of our studies on ligand binding in the class F GPCRs [13, 14, 33], we measured the binding of the only available functional fluorescent WNT – eGFP-tagged mouse WNT-3A [14, 34] - to endogenous FZD_7_ in SW480 cells typifying colorectal cancer, aiming to apply a more relevant cell model. Similar to our recent study [13], we have employed BRET binding assays based on a NanoBiT system (NanoBiT/BRET binding). Here, receptors are N-terminally tagged with the 11 amino acid HiBiT peptide and the subsequent addition of an 18 kDa complementary and cell-impermeable LgBiT protein allows the formation of a stable and catalytically active luciferase - NanoBiT [35]. In the presence of a fluorescent ligand, this setup enables selective analysis of ligand binding to cell surface receptors in living cells and as such can be viewed as an optimal tool to study the association between FZDs and their macromolecular ligands – the WNTs.

In this work, we focus on one FZD paralogue: FZD_7_. We have selected FZD_7_ as a model receptor because of its relevance in cell biology as recently implied by structural studies [36] and drug development campaigns to develop anti-cancer treatments [37]. As such, its function in cancer has been very recently reviewed by Larasati et al. [38]. Among several cancers in which FZD_7_ plays a role, its link to the growth and invasion of colorectal cancer cells has gained considerable attention in the literature [39–42]. FZD_7_ is essential for the maintenance of the intestinal epithelium, involved in cancerogenesis and therefore proposed as a relevant target for the treatment of intestinal cancers [38, 43]. In previous studies, FZD_7_ pharmacology was investigated using primary cells as well as patient-derived immortalised cell lines. To this end, the SW480 colorectal cell model has been used very frequently to assess FZD_7_-mediated effects on e.g. β-catenin-dependent signalling or cell proliferation [40, 44].

Here, we demonstrate that eGFP-WNT-3A interacts with high affinity with endogenous HiBiT-FZD_7_ in SW480 cells. As previously reported for LRP6 (Eckert et al. 2020), we also show that receptor overexpression leads to a significant decrease in ligand binding affinities. Additionally, using the NanoBiT approach we were able to estimate the number of FZD_7_ molecules present at the plasma membrane of SW480 cells. Finally, since the binding experiments were performed in live cells, we have also tested if receptor internalization confounds the observations, and we have demonstrated that agonist-induced internalization did not affect the assessment of ligand affinity. The presented ligand binding studies expand therefore on the current knowledge on WNT-FZD_7_ pharmacology in colorectal cancer. Our findings do not only aid in understanding FZD_7_ biology but can also assist in providing a relevant system for compound screening and validation during the development of FZD_7_-targeting drugs.

## 2. METHODS

### 2.1 Cell culture

HiBiT-FZD_7_ SW480 (generated in this study) and parental SW480 cells (ATCC) were cultured in DMEM supplemented with 10% FBS, 1% penicillin/streptomycin, 1% L-glutamine (complete DMEM; all from Thermo Fisher Scientific) in a humidified CO_2_ incubator at 37°C. All cell culture plastics were from Sarstedt, unless otherwise specified. The absence of mycoplasma contamination was routinely confirmed by PCR using 5′-GGCGAATGGGTGAGTAACACG-3′ and 5′-CGGATAACGCTTGCGACTATG-3′ primers detecting 16S ribosomal RNA of mycoplasma in the media after 2–3 days of cell exposure.

### 2.2 CRISPR-Cas9 genome-editing

SW480 cells expressing genome-edited FZD_7_ with an N-terminal HiBiT tag were generated by CRISPR-Cas9 mediated homology-directed repair (HDR). The sgRNA (5-TCCGTGGTA**CGG**CTGCGCCC-3’; ”-” strand; PAM sequence in bold) used for Cas9 targeting of the N-terminal region of FZD_7_ (NCBI Reference Sequence: NM_003507.2) was designed and synthesised by Integrated DNA Technologies (IDT). The HDR template (5’- GTGCTGGCGCTGCTGGGCGCACTGTCCGCGGGCGCCGGGGCGGTGAGCGGCTGGCGGCTGTTCAAGAAGATTAGCCAGCCGTACCACGGAGAGAAGGGCATCTCCGTGCCGGACC-3’) containing the native FZD_7_ signal peptide and the HiBiT-tag was designed and synthesised by IDT. The addition of silent mutations was not needed for this design because the HDR mutation sufficiently disrupts the guide sequence. SW480 cells at 4 × 10^5^ cells/ml were transfected in suspension using Lipofectamine CRISPRMAX with Plus reagent (Thermo Fisher Scientific) with the following mix: [Cas9] : [HDR template] : [sgRNA] = 10 nM : 3.0 nM : 10 nM, and 1 μM of Alt-R HDR enhancer V2 and seeded in the total volume of 150 μl per well onto a 96-well plate. The cells were cultured for 24 hours in antibiotics-free medium. Subsequently, two consecutive runs of serial dilutions were performed to obtain single cell (clonal) cultures in wells of a 96-well plate. Following a sufficient growth of the cells (approx. 10 weeks), genomic DNA was isolated using the NaOH/Tris-HCl method and, independently, mRNA was isolated with an mRNA Isolation Kit (Roche, # 11 741 985 001) according to the manufacturer’s instructions. mRNA was then used to transcribe cDNA with random primers. To amplify the region encompassing the genome-edited sequence, the following pair of PCR primers was used: 5′- ACCCAGGCTGACGAGTTTTG-3′ and 5′- TAGGGCGCGGTAGGGTAG-3′; predicted product size = 829 bp. The final product size was determined using a Bioanalyzer (Agilent 2100, DNA 1000 Assay). The specific products were validated with Sanger sequencing (Eurofins GATC) with the forward primer 5′- ACCCAGGCTGACGAGTTTTG-3′. Additionally, the protein expression of a CRISPR-Cas9-edited FZD_7_ with an N-terminal HiBiT-tag was assessed with immunoblotting and NanoBiT-emitted luminescence.

### 2.3 Preparation of whole-cell lysates

HiBiT-FZD_7_ SW480 cells were grown to confluency in a 75 cm^2^ cell culture flask. The medium was removed and the cells were washed twice with 5 ml PBS. The HiBiT-FZD_7_ SW480 cells were then detached from the flask using a cell scraper (VWR). The cell suspension in 5 ml PBS was transferred to a 15 ml conical tube and centrifuged once at 400 x g for 5 minutes at room temperature. The supernatant was aspirated and discarded. Cell pellets were stored at -80°C until required or resuspended in an adequate volume of non-phenol red complete DMEM medium. Homogenisation was done on wet ice by sonication (maximum power in 3 x 3 seconds bursts). Next, the samples were centrifuged at 400 x g for 5 min. The supernatants that contained lysed cells were used immediately in experiments.

### 2.4 DNA cloning and mutagenesis

We removed the GS linker adjacent to the HiBiT tag from HiBiT-FZD_7_ [13] using GeneArt site-directed mutagenesis (Thermo Fisher Scientific) and the following primers: 5’-TTCAAGAAGATTAGCCAGCCGTACCACGGA-3’ and 5’-TCCGTGGTACGGCTGGCTAATCTTCTTGAA-3’. The construct was validated by sequencing (Eurofins GATC).

### 2.5 Ligands

Recombinant untagged high-purity human WNT-3A was purchased from RnD Systems (# 5036-WNP-010), dissolved at 200 μg/ml in sterile 0.1% BSA/PBS and stored at 4°C. Recombinant untagged human DKK1 was purchased from RnD Systems (# 5439-DK), dissolved at 100 μg/ml in sterile 0.1% BSA/PBS and stored at 4°C. eGFP-WNT-3A was prepared as described below.

### 2.6 Preparation of eGFP-WNT-3A conditioned media

HEK293F^TM^ suspension cells growing in serum-free Expi293^TM^ expression medium (60 ml, 2.5 × 10^6^ cells/ml) were cotransfected with 10 µg of either pCS2^+^-WNT-3A or pCS2^+^-eGFP-WNT-3A together with 50 µg of pCMV-His-Afamin plasmid using ScreenFect^®^ UP-293 (ScreenFect GmbH) according to the manufacturer’s instructions. The corresponding control conditioned medium was generated from cells transfected with pCS2^+^ plasmid. Cells were first cleared from the HEK293F^TM^ conditioned medium by centrifugation at 310 g for 10 min and then at 3,400 g for 30 min to remove any remaining cellular debris and insoluble material. This “raw” conditioned medium then was concentrated 5-fold using Vivaspin turbo 15 centrifugal concentrators (30,000-molecular-weight-cutoff, Satorius AG, Göttingen, Germany) and exchanged to the desired cell culture medium using Sephadex G-25 PD10 desalting columns (GE Healthcare Bio-Science). The final concentration and integrity of eGFP-WNT-3A in the conditioned medium were determined using both ELISA (GFP ELISA^®^ kit, #ab171581, Abcam) and SDS-PAGE/Western Blot analysis. From the two eGFP-WNT-3A batches (eGFP-WNT-3A batch 1 final concentration: 21 nM, eGFP-WNT-3A batch 2 final concentration: 20 nM), the batch 1 was used in optimization experiments and the batch 2 was used in the experiments presented in this study. The eGFP-WNT-3A concentration was adjusted to 18 nM following the addition of FBS (to 10% final concentration) and HEPES (to 10 mM final concentration; both from Thermo Fisher Scientific). Current WNT purification methods allow only limited WNT concentration to be obtained from a conditioned medium [45]. For the validation of the eGFP-WNT-3A batches please see **Figure S1**.

### 2.7 LgBiT/HiBiT calibration curve

One day before the experiment, SW480 cells were seeded at a density of 4 × 10^4^ cells/well (100 μl) onto a poly-D-lysine-coated (Thermo Fisher Scientific) black 96-well cell culture plate with a transparent flat bottom (Greiner BioOne). A white adhesive tape (VWR) was attached to the bottom of the plate before the measurement. On the day of the experiment, cells were washed once with 200 µl Hanks’ balanced salt solution (HBSS, HyClone) and maintained in 80 µl of complete, non-phenol red DMEM (HyClone) supplemented with 10 mM HEPES (HyClone). HiBiT control protein (#N301A, Promega) dilutions were prepared in the same medium and added (10 µl/well). Next, 10 µl of a mix of LgBiT (1:20 dilution; #N2421, Promega) and furimazine (1:10 dilution; #N2421, Promega), prepared in the above-described medium, were added. After an incubation period of 10 min at 37°C (no CO_2_), luminescence (460-500 nm, 200 ms integration time) was recorded using a TECAN Spark microplate reader (TECAN). To estimate the number of FZD_7_ molecules on the cell surface, we used the following: the formula from the calibration curve, the Avogadro constant = 6,02214076 × 10^23^ moles^-1^, average number of cells/well = 72500 and solution volume in a well = 100 μl.

### 2.8 NanoBiT/BRET binding assay

HiBiT-FZD_7_ SW480 cells were left untransfected or transiently transfected in suspension using Lipofectamine^®^ 2000 (Thermo Fisher Scientific). A total of ca. 4 × 10^5^ cells were transfected in 1 ml with 100 ng of HiBiT-FZD_7_ plasmid DNA and 900 ng of pcDNA3.1 plasmid DNA (low overexpression) or 1000 ng of HiBiT-FZD_7_ plasmid DNA (high overexpression). The cells (100 µl) were seeded onto a poly-D-lysine-coated black 96-well cell culture plate with a transparent flat bottom (Greiner BioOne). Before BRET measurements, a white adhesive tape (VWR) was attached to the plate bottom. Twenty-four hours post-transfection, the cells nearly doubled (average total cell number in a well = 72500). They were washed once with 200 µl of HBSS (HyClone). In the kinetic binding experiments, the cells were preincubated with 50 µl of a mix of Nluc substrate endurazine (1:50 dilution; #N2571, Promega) and LgBiT (1:50 dilution; #N2421, Promega) in a complete, non-phenol red DMEM (HyClone) supplemented with 10 mM HEPES (HyClone) for 90 min at 37 °C without CO_2_. Subsequently, 50 µl of eGFP-WNT-3A conditioned medium or control medium supplemented with 10% FBS and 10 mM HEPES were added, and the BRET signal was measured every 87 s for 300 min at 37 °C. In the saturation-binding experiments, the cells were incubated with different concentrations of eGFP-WNT-3A conditioned medium (90 µl) supplemented with 10% FBS and 10 mM HEPES for 180 min at 37°C (no CO_2_). Next, for saturation binding experiments, 10 μl of a mix of furimazine (1:10 dilution; #N2421, Promega) and LgBiT (1:20 dilution; #N2421, Promega) were added. The cells were incubated for another 10 min at 37 °C (no CO_2_) before the BRET measurements. The BRET ratio was defined as the ratio of light emitted by eGFP-WNT3A (energy acceptor) and light emitted by HiBiT-FZD_7_ (energy donor). Net BRET ratio was defined as BRET ratio _ligand-treated wells_ – BRET ratio _vehicle-treated wells_. The BRET acceptor (520–560 nm) and BRET donor (460–500 nm) emission signals were recorded with 200 ms integration time using a TECAN Spark microplate reader (TECAN). Cell surface expression of HiBiT-FZD_7_ was assessed by measuring luminescence of vehicle-treated wells (no BRET acceptor) in the NanoBiT/BRET binding assays.

### 2.9 Receptor internalization

HiBiT-FZD_7_ SW480 cells (4 × 10^4^ cells in 100 μl) were seeded onto a well of a poly-D-lysine-coated black 96-well cell culture plate with a transparent flat bottom (Greiner BioOne). Twenty-four hours later, the cells were washed once with 200 µl of HBSS (HyClone). Next, 45 μl of a complete, non-phenol red DMEM (HyClone) supplemented with 10 mM HEPES was added to the wells followed by the addition of 45 µl of 18 nM (9 nM final concentration) of high-purity recombinant untagged WNT-3A. The ligand was added every 30 min for 210 min, keeping the cells at 37 °C without CO_2_. Finally, 10 μl of a mix of furimazine (1:10 dilution; #N2421, Promega) and LgBiT (1:20 dilution; #N2421, Promega) was added 20 min after the last WNT-3A addition and the plate was incubated for 10 min at 37 °C without CO_2_. Before luminescence measurements, the white adhesive tape was attached to the plate bottom. Next, NanoBiT emission (460–500 nm, 200 ms integration time) was measured using a TECAN Spark microplate reader (TECAN).

### 2.10 Immunoblotting

Whole-cell lysates were obtained using Laemmli buffer with 0.5% NP-40 and 5% β-mercaptoethanol. Lysates were sonicated and analyzed on 4–20 % Mini-PROTEAN TGX precast polyacrylamide gels (Bio-Rad) and transferred to PVDF membranes using the Trans-Blot Turbo system (Bio-Rad). After blocking with 5% milk in Tris/NaCl/Tween20 buffer (TBS-T), membranes were incubated with the following primary antibodies in the blocking buffer: rabbit anti-GAPDH (1:2,500; Cell Signaling Technology #2118) and mouse anti-HiBiT (1.0 µg/ml; Promega clone 30E5), overnight at 4 °C. After multiple washing steps with TBS-T, membranes were incubated with horseradish peroxidase-conjugated secondary antibodies in the blocking buffer for 1 h at room temperature (1:3,000 rabbit anti-goat, Sigma-Aldrich #A5420 or 1:3,000 goat anti-mouse, Thermo Fisher Scientific #31430). After multiple washing steps, membranes were incubated for 2 minutes with Clarity Western ECL Blotting Substrate (Bio-Rad) and proteins visualized on ChemiDoc (Bio-Rad). The uncropped blots can be found in the **Figure S2**.

### 2.11 Data and statistical analysis

All data were analyzed in GraphPad Prism 8 using built-in equations. All data presented in this study come from at least five individual experiments (biological replicates) with each individual experiment performed typically in duplicates (technical replicates) for each tested concentration/condition, unless otherwise specified in a figure legend. Data points on the binding curves represent mean ± SEM. Saturation binding data were fit using a one-site specific model. Kinetic binding data were analyzed using the association model with two or more hot ligand concentrations. Equilibrium dissociation constant values (*K*_d_) representing ligand binding affinities are reported as a best-fit *K*_d_ with SEM. *K*_d_ values were compared using an extra-sum-of-square F-test (P<0.05). WNT-3A-induced HiBiT-FZD_7_ internalization data were fitted to a one-phase decay equation, normalized to the vehicle-treated wells (100%) and presented with 95% confidence intervals (95% CI). HiBiT-LgBiT calibration data were analyzed using a simple linear regression model with luminescence intensities and HiBiT concentrations presented on logarithmic scales. Cell surface expression data in the **Figure 2B** and **Figure S4** are presented as mean ± SEM and were analyzed for differences with one-way analysis of variance (ANOVA) followed by Tukey’s test; **** P ≤ 0.0001, * P ≤ 0.05.

## 3. RESULTS

### Genome engineering of SW480 cells

We generated a genome-edited SW480 clonal cell line expressing FZD_7_ tagged on the N-terminus with the 11-amino acid HiBiT-tag (hereafter referred to as HiBiT-FZD_7_ SW480 cells). Sequencing of the FZD_7_-specific PCR product amplified from genomic DNA resulted in one distinct sequence, implying that a homozygous clonal line was obtained (**Figure S3A**). Next, these results were confirmed with the sequencing of the FZD_7_-specific PCR product amplified from cDNA reversely-transcribed from total cellular mRNA which also presented itself with a single distinct sequence indicating the presence of RNA coding for HiBiT-FZD_7_ and not for an alternative, untagged FZD_7_ (**Figure S3B**). In agreement with the sequencing data, we detected the expression of full-length HiBiT-FZD_7_ by immunoblotting (**Figure 1A**). Additionally, we detected bioluminescence following incubation of the live cells with furimazine and the membrane-impermeable LgBiT, indicative of cell-membrane localization of the tagged protein in the genome-edited SW480 cells (**Figure 1B**).

**Fig. 1:**
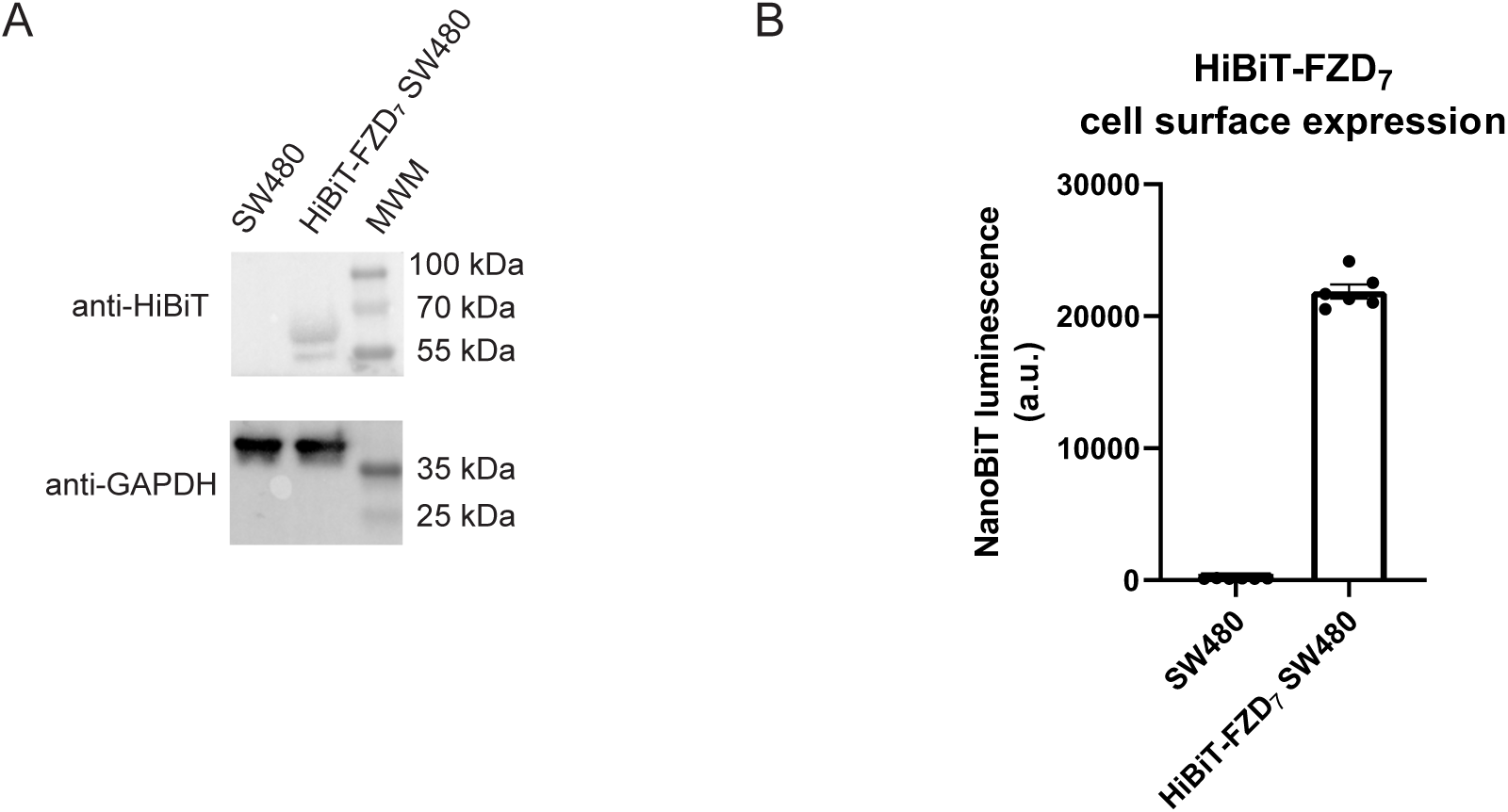
Validation of HiBiT-FZD_7_ SW480 cells. **A.** Validation of cellular expression of endogenous HiBiT-FZD_7_ in parental and CRISPR-Cas9 genome-edited SW480 cells. Cell lysates were analyzed by immunoblotting using anti-HiBiT antibody and anti-GAPDH served as a loading control. The predicted molecular weight of HiBiT-FZD_7_=65.0 kDa. Western blot experiments were repeated five times with similar results. MWM – molecular weight marker. **B.** Surface expression of HiBiT-FZD_7_ was quantified by measuring NanoBiT-emitted luminescence. Raw data are shown from n = 6 individual experiments and are presented as mean ± SEM.

### Quantification of FZD_7_ expression in SW480 cells

Expression levels of HiBiT-FZD_7_ molecules at the plasma membrane of live SW480 cells were estimated using NanoBiT-emitted luminescence and a LgBiT-HiBiT calibration curve. Here, the addition of increasing concentrations of purified HiBiT in the presence of fixed working dilutions of LgBiT and furimazine led to a linear increase in luminescence presented on a logarithmic scale (**Figure 2A**). In this manner, we could estimate the number of HiBiT-FZD_7_ molecules present on the surface of a single cell (**Figure 2B**). Thus, a SW480 cell expresses on average 1373 ± 349 FZD_7_ proteins on its surface. Similarly, we applied these calculations also to overexpressed and highly overexpressed conditions (**Figure 2B**), which resulted in average values of 7034 ± 806 and 42609 ± 2383 membrane receptors/cell, respectively.

**Fig. 2:**
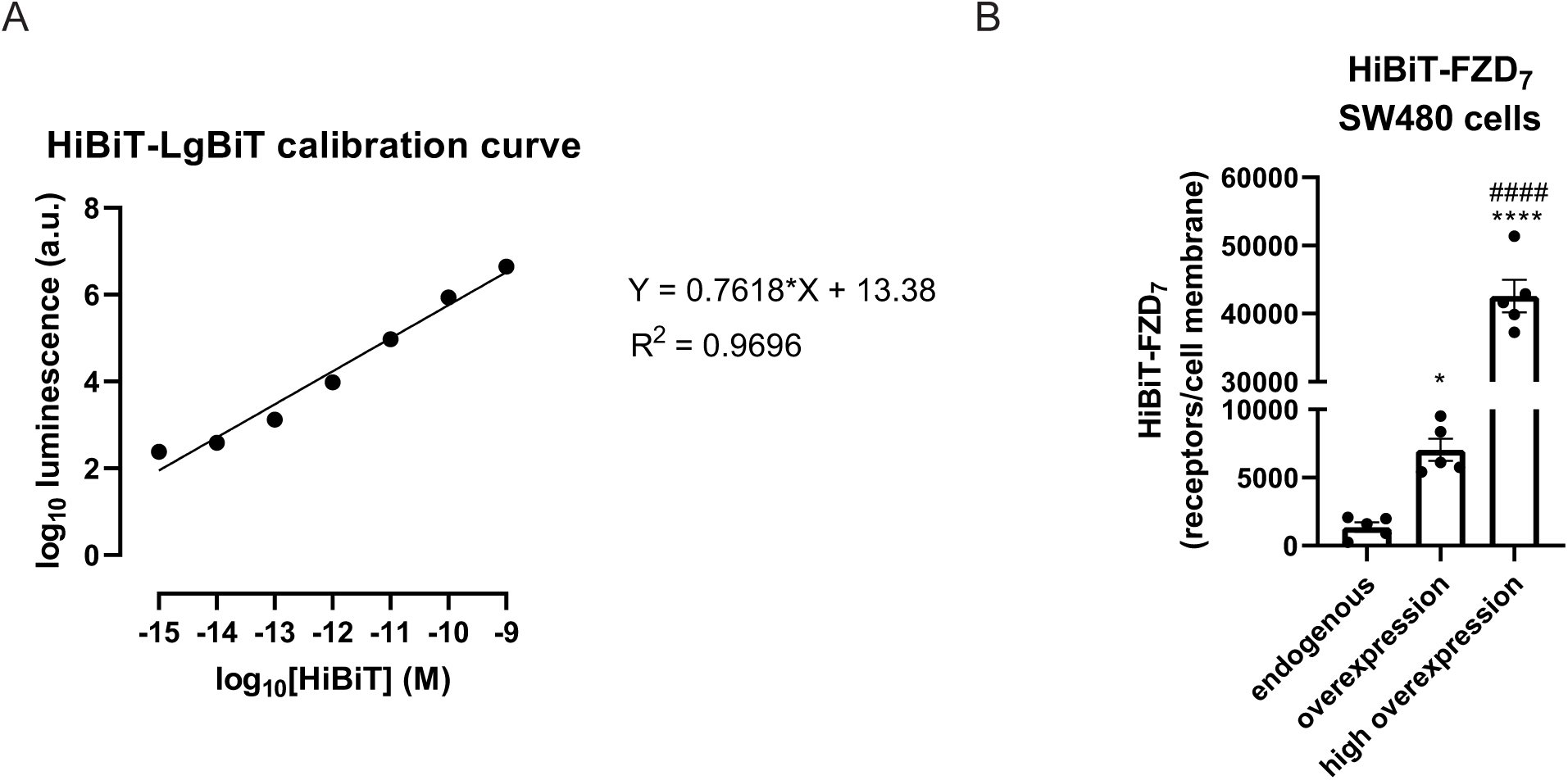
Estimation of HiBiT-FZD_7_ expression on the cell surface of HiBiT-FZD_7_ SW480 cells. **A.** Concentration-dependent effect of purified HiBiT on complemented luminescence following incubation with 1:200 LgBiT and 1:100 furimazine. SW480 cells were plated onto a black transparent-bottom 96-well plate and luminescence was measured in 100 μl of a complete non-phenol red DMEM. Raw data are shown from n = 5 individual experiments and are presented as mean ± SEM; a.u. -an arbitrary unit. **B.** Surface expression of HiBiT-FZD_7_ under endogenous, overexpressed and highly overexpressed conditions in the CRISPR-Cas9 genome-edited HiBiT-FZD_7_ SW480 cells was measured by quantifying NanoBiT-emitted luminescence and the number of membranous receptor molecules estimated using the equation from the panel A. The average number of cells (72500 cells/well) was used in the calculations. Data are shown from n = 5 individual experiments and are presented as mean ± SEM; * denotes comparison with the endogenous expression levels, # denotes comparison with the overexpressed levels. Please refer to the **Fig. S4** for the raw luminescence data.

### Kinetic binding of eGFP-WNT-3A to endogenous and overexpressed HiBiT-FZD_7_

HiBiT-FZD_7_ SW480 cells were used to study the binding kinetics of eGFP-WNT-3A to endogenous FZD_7_ as well as to FZD_7_ overexpressed at two different levels. For these experiments, cells were first incubated with the complementary LgBiT protein and the long-lasting luciferase substrate endurazine. After 90 minutes, eGFP-WNT-3A was added to final concentrations of either 0.9 nM, 1.8 nM or 2.7 nM and BRET readings were recorded over a five-hour period at 37°C (**Figure 3**). This kinetic analysis showed that eGFP-WNT-3A associates to endogenously expressed HiBiT-FZD_7_ in a rapid and saturable manner, reaching a plateau after approximately 80 minutes. However, in the case of receptor overexpression, the eGFP-WNT-3A binding occurred with a slower on-rate to reach only nearly saturable levels after the five hours of experimental duration (**Table 1**).

**Fig. 3:**
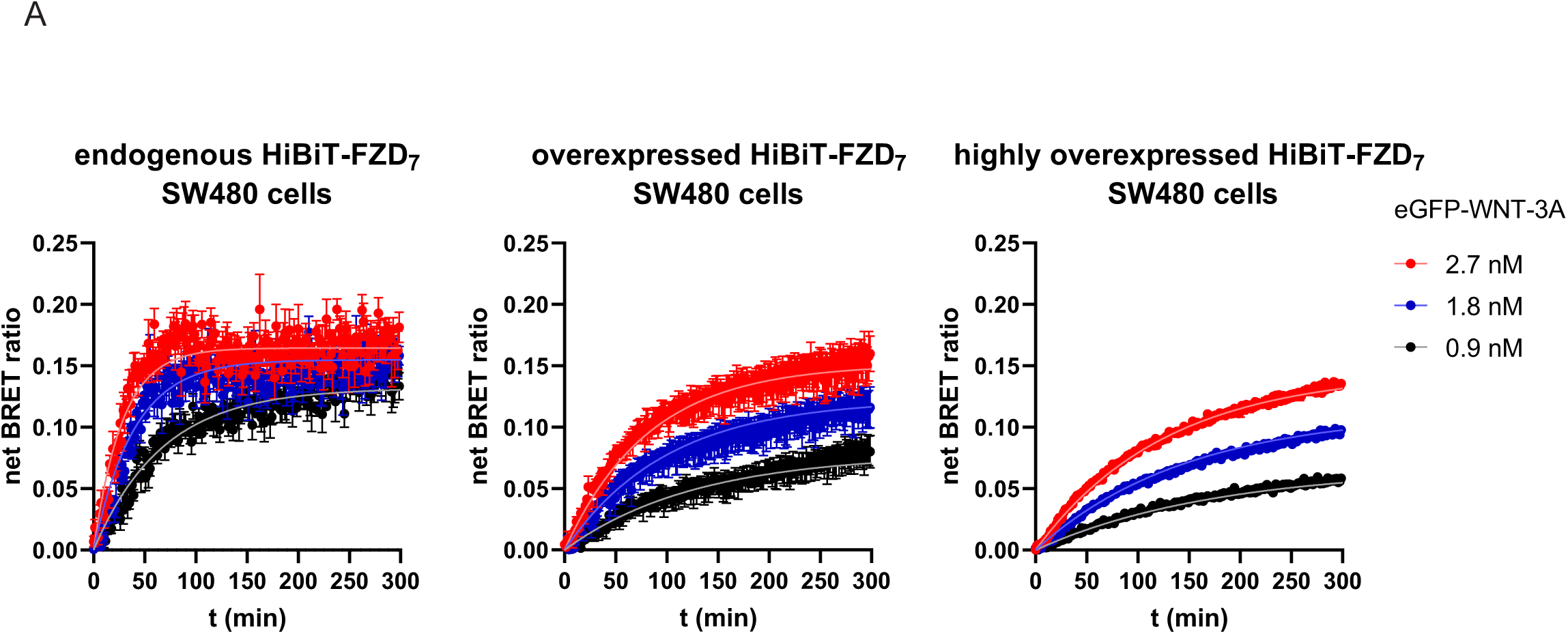
eGFP-WNT-3A binding kinetics. Association kinetics of eGFP-WNT-3A to HiBiT-FZD_7_ expressed at the three different levels were determined using 0.9, 1.8 and 2.7 nM by detection of NanoBiT/BRET in living CRISPR-Cas9 genome-edited HiBiT-FZD_7_ SW480 cells. NanoBiT/BRET was sampled once every 87 seconds for 300 minutes. Raw data were fitted to the ”two or more hot concentrations model” and are presented as mean ± S.E.M. from n = 5 individual experiments. Kinetic parameters are summarized in **Table 1**.

**Table 1:**
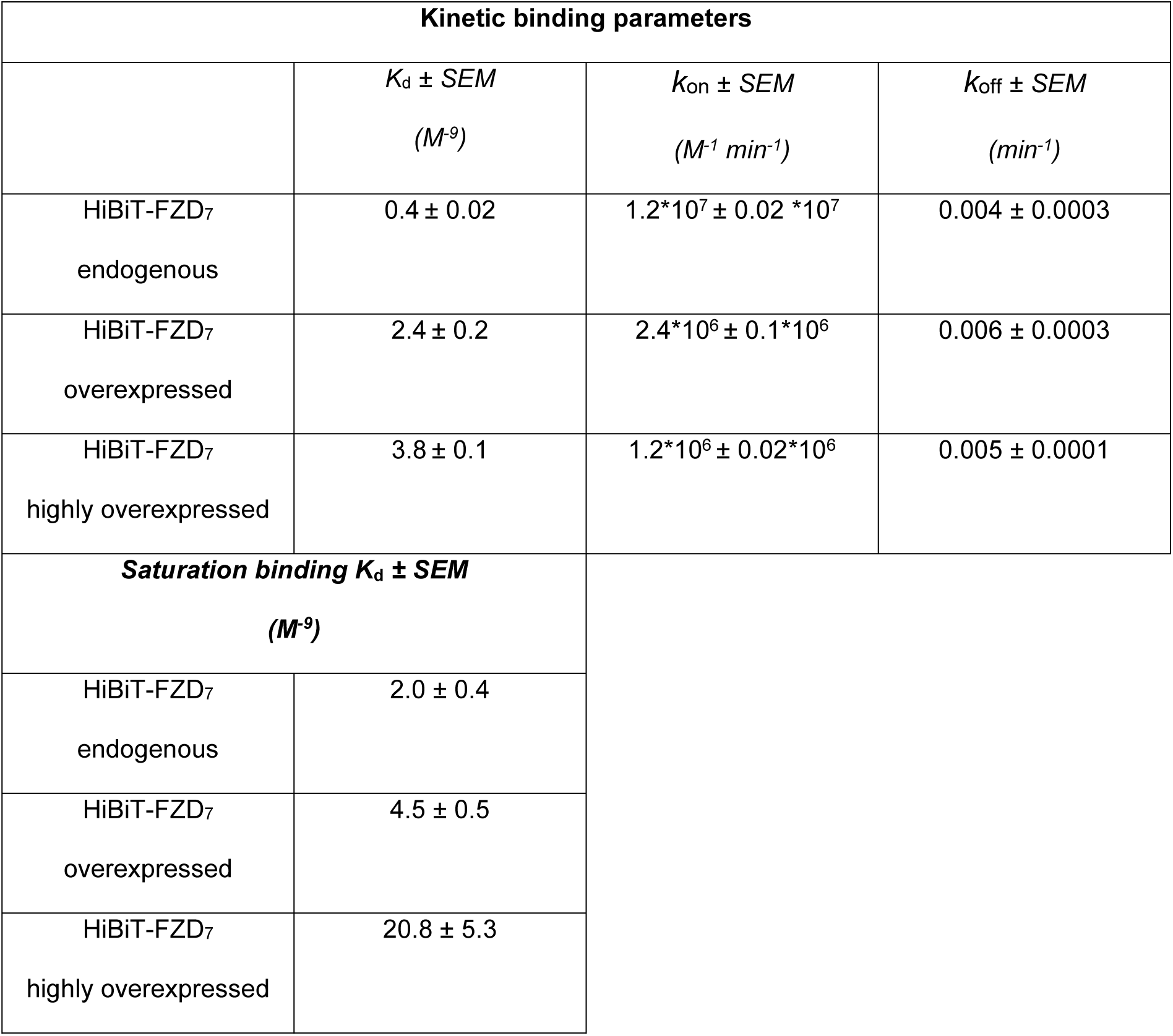
Kinetic binding and saturation binding parameters of eGFP-WNT-3A binding to HiBiT-FZD_7_. Values are based on data from n=5 individual experiments (shown in **Figure 3** and **Figure 4**) and presented as a best-fit value ± SEM.

### eGFP-WNT-3A saturation binding to endogenous and overexpressed HiBiT-FZD_7_

To define the saturation binding affinity of eGFP-WNT-3A, we incubated HiBiT-FZD_7_ SW480 cells with a full concentration range (0.027 nM to 9.0 nM) of eGFP-WNT-3A (**Figure 4**). The net BRET ratio representing ligand-receptor binding increased in a clear concentration-dependent manner and reached saturation for SW480 cells expressing HiBiT-FZD_7_ at endogenous and low overexpression levels (**Figure 4** **top left** and **top right**). On the contrary, eGFP-WNT-3A binding to HiBiT-FZD_7_ overexpressed at high levels did not reach maximal asymptotic values (**Figure 4** **bottom**). The apparent affinities of eGFP-WNT-3A/FZD_7_ interactions were determined using a one-site specific non-linear regression model and the *K*_d_ values are shown in **Table 1**. We have used the data from individual experiments underlying the results presented in the **Figures 2** and **4** to display the correlation between HiBiT-FZD_7_ levels and eGFP-WNT-3A/FZD_7_ binding affinities. These data are presented in the **Figure 5**.

**Fig. 4:**
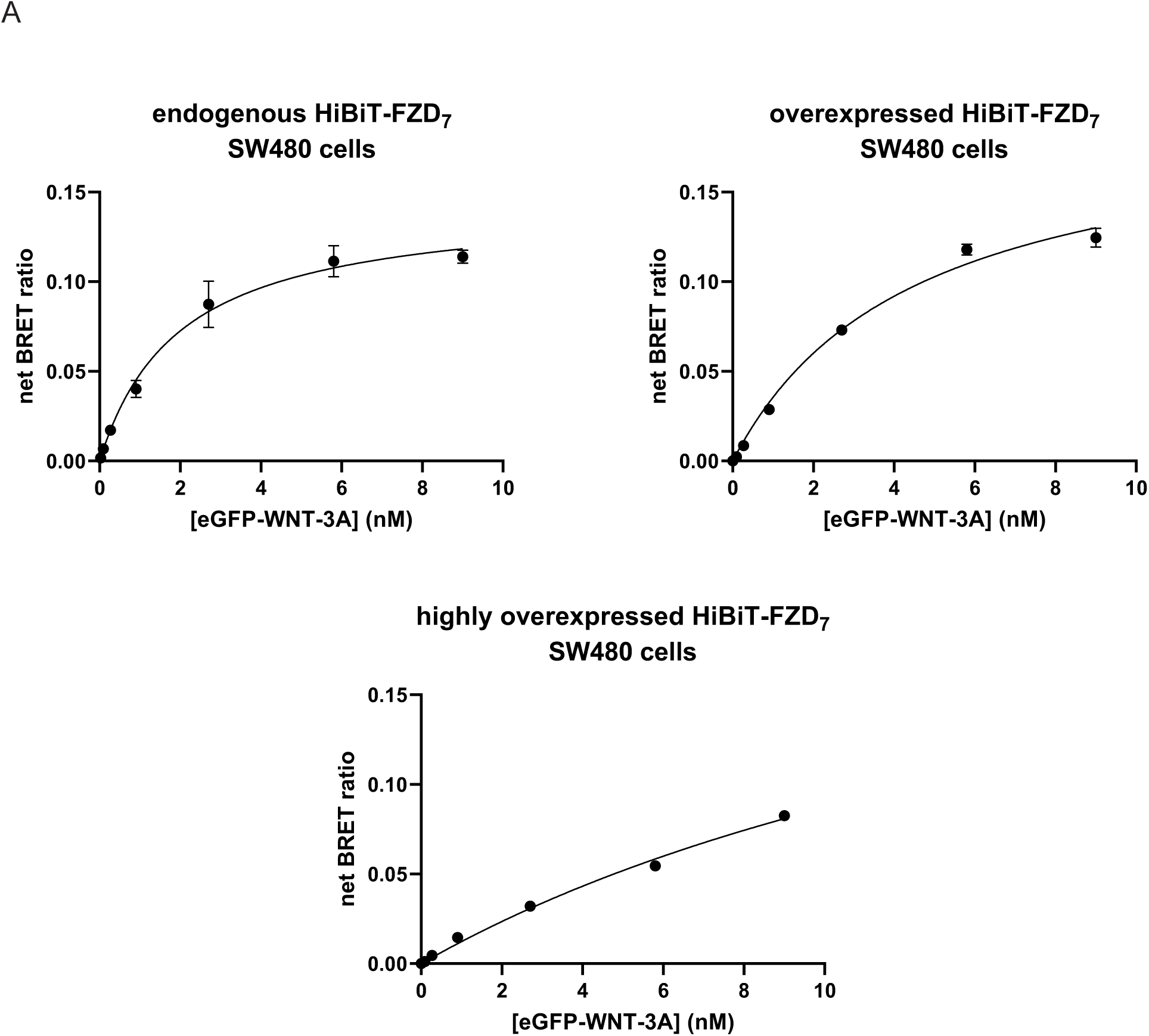
eGFP-WNT-3A saturation binding. Saturation binding of the eGFP-WNT-3A to HiBiT-FZD_7_ expressed at the three different levels in living CRISPR-Cas9 genome-edited HiBiT-FZD_7_ SW480 cells were determined using a full concentrations range (0.027 nM to 9 nM) following 180 min incubation. Raw data are presented as means ± SEM from n = 5 individual experiments, fitting a one-site specific model. The equilibrium dissociation constants (*K*_d_) are summarized in **Table 1**.

**Fig. 5:**
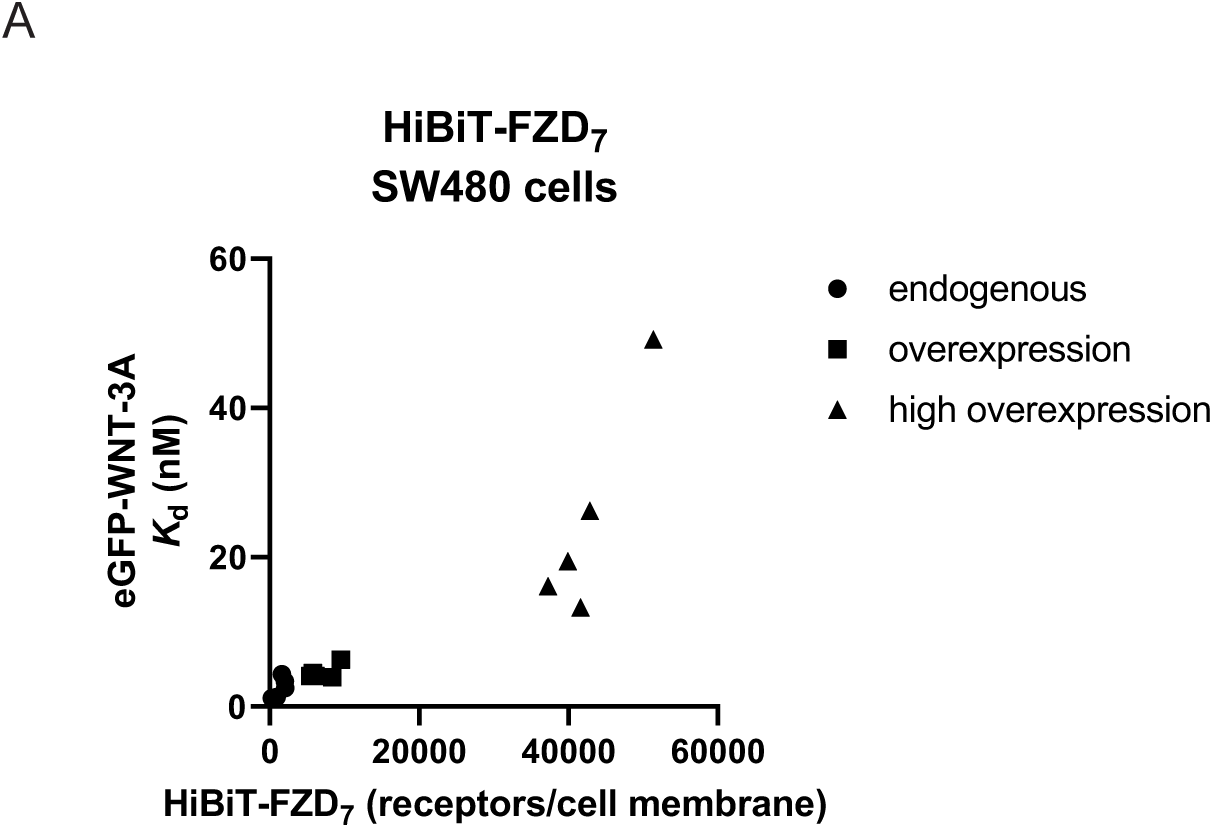
eGFP-WNT-3A binding affinity decreases with increasing numbers of HiBiT-FZD_7_. The graphs present a correlation between eGFP-WNT-3A *K*_d_ in nM (left axis) and HiBiT-FZD_7_ expression in molecules/cell membrane. Data points represent *K*_d_ and receptor molecule numbers come from each of 5 individual experiments for each condition (15 data points in total).

### WNT-3A-induced HiBIT-FZD_7_ internalization and its effect on ligand binding measurements

Since WNT-3A is a FZD agonist, we aimed to determine the effects of a potential WNT-3A-induced HiBiT-FZD_7_ internalization on the assessment of ligand affinity by measuring NanoBiT-emitted luminescence following the stimulation of HiBiT-FZD_7_ SW480 cells with WNT-3A. We used a recombinant untagged, high-purity, human WNT-3A at a concentration of 9 nM, matching the highest fluorescent ligand concentration used in the ligand binding assays. Incubation with a fixed WNT-3A concentration led to a time-dependent decrease in luminescence emitted by NanoBiT (**Figure 6A**). This decrease – indicative of HiBiT-FZD_7_ internalization – with half-life (*t*_1/2_) of 34.2 min (95% CI = 21.0 - 60.5) plateaued after an incubation period of 65 min (plateau = 64.7 min; 95% CI = 58.1 - 69.1). From this time point on, approximately 60-70% of the initial receptor pool was still present at the cell membrane. Next, to measure the impact of receptor internalization on the ligand binding affinity measurements, we did not use live cells but instead we incubated internalization-incapable HiBiT-FZD_7_ SW480 cell lysates (note: endogenous expression levels) with increasing concentrations of eGFP-WNT-3A. We used the same setup as in the saturation binding on live cells. Again, we detected a concentration-dependent saturable binding of eGFP-WNT-3A to HiBiT-FZD_7_ (**Figure 6B**) that presented itself with a *K*_d_ value of 2.5 ± 0.7 nM, which was not statistically different from the saturation binding *K*_d_ obtained with live cells (please see the **Table 1**).

**Fig. 6:**
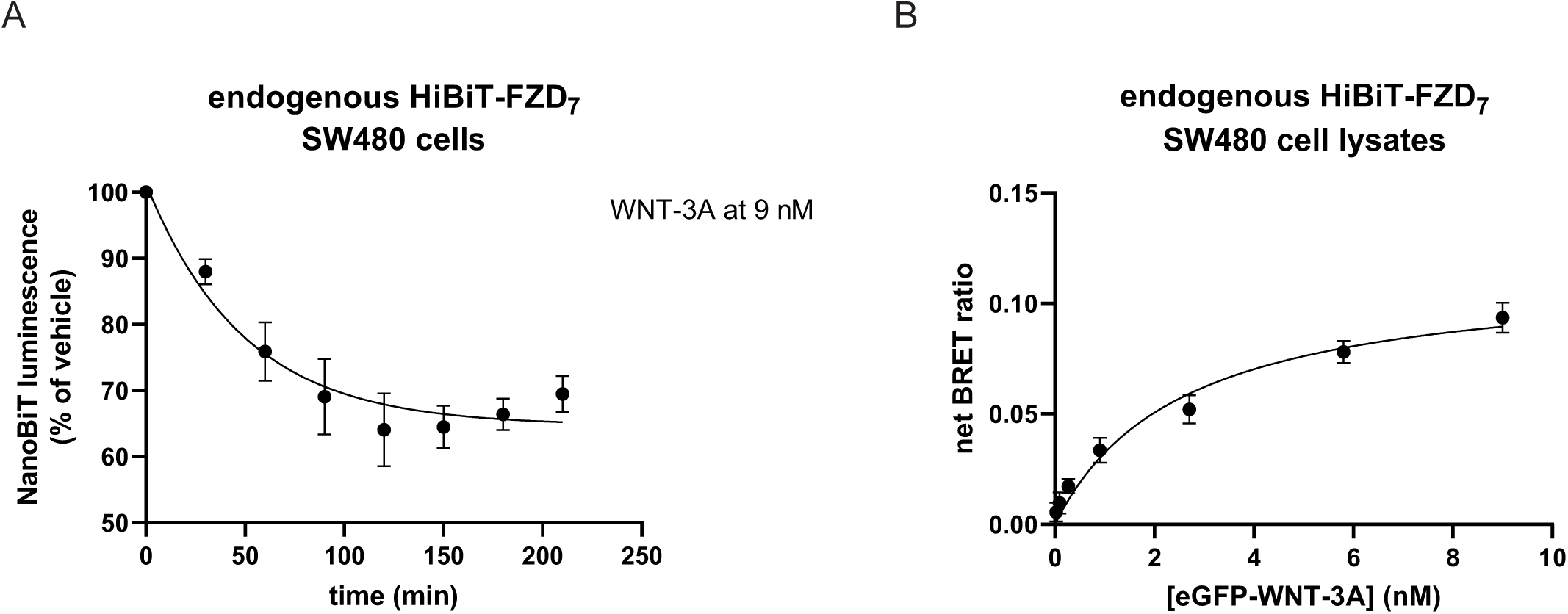
WNT-3A-induced internalization of HiBiT-FZD_7_ does not affect NanoBiT/BRET binding measurements. **A.** Time course of recombinant untagged high purity human WNT-3A-induced internalization of endogenous HiBiT-FZD_7_ in HiBiT-FZD_7_ SW480 cells. Luminescence of cell surface complemented HiBiT–LgBiT (NanoBiT) was measured after 210 min following additions of 9 nM WNT-3A in 30 min intervals. Normalized data are presented as mean ± SEM from n = 5 individual experiments and fitted to a one-phase decay model. **B.** Saturation binding of the eGFP-WNT-3A to endogenous HiBiT-FZD_7_ in whole-cell lysates of CRISPR-Cas9 genome-edited HiBiT-FZD_7_ SW480 were determined using a full concentrations range (0.027 nM to 9 nM) following 180 min incubation. Raw data are presented as means ± SEM from n = 5 individual experiments, fitting a one-site specific model.

## 4. DISCUSSION AND CONCLUSIONS

Here, we have used NanoBiT/BRET and CRISPR-Cas9 genome editing to compare eGFP-WNT-3A binding to HiBiT-FZD_7_ expressed either from its endogenous promotor or from an overexpressing CMV promoter in SW480 cells typifying colorectal cancer. Additionally, we have estimated the number of the receptor molecules on the cell surface of these cells and assessed WNT-3A-induced internalization of HiBiT-FZD_7_ by measuring NanoBiT-emitted luminescence. This is the first study to quantify WNT association with a tagged FZD expressed from its endogenous promoter. Below we will discuss the advances of this work, the interpretation of the data and the impact this work will have on our ongoing efforts to study and quantify WNT-FZD interactions. Nonetheless, it has to be noted that there have only been two studies on WNT-FZD BRET binding so far [13, 14] and hence we are mostly limited to discussing our previous findings or linking our results to the data from the class A GPCRs[32].

We performed the experiments in SW480 cells to provide a more relevant model to study WNT-FZD biology than the commonly used HEK293 cells, which also provided an opportunity to compare WNT binding to overexpressed FZDs with binding to FZDs expressed at lower, endogenous levels [13]. We used CRISPR-Cas9 genome editing in SW480 cells to insert an 11-amino acid HiBiT tag to the 5’ end of the endogenous FZD_7_ locus. We have previously demonstrated that addition of a HiBiT tag does not compromise receptor’s functions as recombinant overexpressed human HiBiT-FZD_7_ was capable of binding eGFP-WNT-3A and mediating WNT-3A-induced β-catenin-dependent signaling [13]. It has to be noted that the addition of the tag was the only genome edition of the FZD_7_ gene we performed here as we have kept the native FZD_7_ signal peptide and we have not introduced any silent mutations or restriction sites. Here, it should be mentioned that SW480 cells are defined as hyperdiploid [46]. As such, non-diploid cells have posed difficulties in establishing homozygous clones following genome editing with CRISPR-Cas9 [30]. Hence, to analyze the efficiency of CRISPR-Cas9 tagging, and to prove the homozygotic status of the clones, we performed Sanger sequencing. Sequencing of the FZD_7_-specific PCR product amplified from genomic DNA resulted in a single distinct peak, indicating that we indeed generated a homozygous clonal line. These results were then further supported by sequencing of an 829 bp product from the mRNA template where again only a single sequencing signal was visible indicative of translation of one FZD_7_ variant – HiBiT-FZD_7_. Finally, the correct size and proper cell-surface trafficking of this membrane receptor have been confirmed with immunoblotting using anti-HiBiT antibody and NanoBiT-emitted luminescence, respectively. In summary, our analysis validated the generation of engineered homozygous SW480 cell system.

Next, we have estimated the number of FZD_7_ molecules on the cell surface of the HiBiT-FZD_7_ SW480 cells using NanoBiT-emitted luminescence. We have performed assays in three experimental setups in the HiBiT-FZD_7_ SW480 cells, using 1) untransfected, 2) FZD_7_-overexpressing or 3) FZD_7_-highly-overexpressing cells. Cell-surface expression of HiBiT-FZD_7_ has increased approx. 5-fold following transfection of the cells with HiBiT-FZD_7_ plasmid (10 ng of receptor plasmid DNA in a well). In the case of the high overexpression setup (100 ng of receptor plasmid DNA in a well), the FZD_7_ levels were even boosted as much as 31 times. These cells have subsequently been used to measure eGFP-WNT-3A binding to HiBiT-FZD_7_. First, we performed NanoBiT/BRET binding in a kinetic format. We monitored the association of the fluorescently tagged WNT to HiBiT-tagged FZD_7_ over a five-hour time period sampling BRET. We have detected clear receptor expression-dependent changes in the kinetic parameters of binding, especially in the association rate constant *k*_on_. To this end, eGFP-WNT-3A binding to endogenous HiBiT-FZD_7_ saturated following approximately 80 min of incubation for two of three ligand concentrations. On the contrary, the association to the overexpressed and highly overexpressed receptors presented itself with nearly-saturable curves, which did not reach full saturation during the experiment time. Thus, we observed changes in the kinetic binding parameters with particularly dramatic differences in the association rate constants (*k*_on_) leading to differences in kinetic *K*_d_ values. Even though the luminescence values (**Figure S4**) for endogenously expressing cells were close to the lower detection limit of the plate reader, we were still able to detect eGFP-WNT-3A binding to HiBiT-FZD_7_ in a robust manner. We also performed saturation binding studies, in which we incubated the HiBiT-FZD_7_ cells with the fluorescent ligand. It needs to be noted that incubation time between three to five half-times for dissocation (*t*_1/2_=ln2/*k*_off_) is recommended to reach equilibrium in saturation binding experiments [47, 48]. For our experiments, we would have to use impractical incubation times between 9-14 hours. Instead, we followed effective equilibration times for high affinity ligands as discussed by Hoare [48]. Thus, we incubated eGFP-WNT-3A with the cells for three hours to subsequently report apparent binding affinities that are likely underestimates of true binding affinities reported in the kinetic binding study. With this caveat, the data also support the notion that higher receptor levels of HiBiT-FZD_7_ lead to a lower apparent binding affinity (higher apparent *K*_d_ in nM) of eGFP-WNT-3A. Importantly, the apparent binding affinities of eGFP-WNT-3A to highly overexpressed HiBiT-FZD_7_ in SW480 cells were comparable to those we have obtained in HEK293A cells under similar experimental conditions [13].

In classical radioligand binding assays using cell membranes or whole-cell lysates, an increase in receptor expression could lead to ligand depletion due to the receptor binding, partition of the tracer into cell membranes and other non-specific interactions [49]. To prevent this, the receptor concentration should generally not exceed a tenth of *K*_d_ [47]. However, problems with applying classical ligand binding concepts to novel methodologies and microplate format have been recently discussed [48]. Still, having used the formulas to calculate ligand depletion that were supplied with that study, we argue that ligand depletion does not impact our analysis. Furthermore, selective targeting of membrane receptors in the NanoBiT/BRET setup greatly limits any impact of non-specific interactions and compartmentalization that cannot interfere with the BRET readout due to the far distance from the NanoBiT-tagged receptor at which they occur. Therefore, we speculate that the observed increase in ligand-receptor affinity for WNT-3A/FZD_7_ with reduced receptor expression is not the result of a sub-optimal technical setup. Along these lines, recent studies on NanoBRET binding to Class A GPCRs have generally reported on a similar tendency without, however, discussing potential causes in detail [32, 50]. Similarly, for the WNT receptor LRP6, an increase in the number of LRP6-mCherry molecules at the cell surface led to a decrease in DKK1-eGFP binding affinity measured with line-scanning fluorescence correlation spectroscopy (lsFCS) in HEK293 cells and NCI-H1730 lung cancer cells [26]. Indeed, LRP6-mCherry expressed from endogenously-tagged (CRISPR-Cas9-edited) H1703 cells, or a stable H1703 cell line selected for low expression, revealed a three-fold increase in binding affinities compared to transient overexpression of LRP6-mCherry in the same cells [26]. This affinity modulation was linked to a potential role of receptor-receptor interactions that become more prevalent following receptor overexpression. Along these lines, FZD_7_ can form homodimers upon overexpression [51]. Additionally, WNT-3A binds to both FZD_7_ and LRP6 to initiate β-catenin-dependent signalling [52]. To assess the contribution of eGFP-WNT-3A binding to LRP5/6 to the observed ligand affinity for HiBiT-FZD_7_, we used the saturation binding setup to measure eGFP-WNT-3A association with endogenous HiBiT-FZD_7_ in the presence of 11.6 nM (300 ng/ml) of unlabelled recombinant human DKK1 (**Figure S5**), which blocks the endogenous LRP5/6. In this way, we created conditions where available HiBiT-FZD_7_ binding sites for eGFP-WNT3A likely outnumbered available LRP5/6 binding sites. However, in these saturation binding experiments in the presence of DKK1, we did not detect a statistically significant difference in the *K*_d_ of eGFP-WNT-3A to endogenous HiBiT-FZD_7_ in the presence of DKK1 compared with the results presented in **Table 1** (*K*_d_ with DKK1 = 2.4 ± 0.3 nM) indicating that LRP5/6 binding does not contribute to the WNT affinity to HiBiT-FZD_7_ expressed in a native cell context. All in all, our data demonstrate that higher FZD_7_ surface expression levels result in a decrease in WNT-3A binding affinity in a colorectal cancer model. However, further work is required to understand how the cellular context governs these various ligand binding affinities to receptors present in different numbers on the surface of SW480 cells.

Finally, as the fluorescent probe used in this study is a proteinaceous agonist, we aimed to assess potential WNT-3A-induced HiBiT-FZD_7_ internalization and verify its impact on the NanoBiT/BRET measurements of binding affinity. As such, this FZD paralogue has been shown to possess a dynamic membrane expression profile in Wilms’ tumour as well as to internalize upon incubation with FZD_7_-specific Fab in human embryonic stem cells [53, 54]. Here, we assessed the internalization of endogenous HiBiT-FZD_7_ upon stimulation with recombinant human high-purity WNT-3A by measuring NanoBiT-emitted luminescence in a time-dependent manner. We could measure a gradual decrease in the luminescence signal, indicative of a reduction in the number of the membrane-bound HiBiT-FZD_7._ Subsequently, to assess whether the NanoBiT/BRET measurements of eGFP-WNT-3A saturation binding to HiBiT-FZD_7_ in live cells are compromised by the steady reduction in the pool of membranous receptors, we ran the saturation binding in whole-cell lysates. These experiments did not show differences in *K*_d_ hence indicating that binding measurements are likely not subject to errors due to HiBiT-FZD_7_ internalization. Moreover, an energy transfer from a NanoBiT-tagged receptor to a receptor-bound fluorescent ligand could most likely still be detected from an internalized receptor-ligand complex.

In summary, we have shown that eGFP-WNT-3A binding affinity to HiBiT-FZD_7_ in SW480 cells decreases with an increase in receptor concentration. This study supports the need of studying WNT-FZD interactions at low levels of FZD expression and provides - at least in part - an explanation for the requirement of high WNT concentrations for functional readouts in FZD overexpression systems [18, 19, 21]. Interestingly, it has still not been possible to determine endogenous WNT concentrations in tissues or tumours relative to the FZD concentration. Biologically, our findings argue that slight WNT overexpression for example in the tumour microenvironment indeed can result in a substantial signal input provided that the change of ligand concentration is around the *K*_d_ of the receptor.

With the development of other fluorescent WNTs, FZD-binding small molecules and the generation of relevant cell lines expressing low levels of FZD, our concepts and tools can be used to more accurately compare WNT-FZD selectivity and optimize drug development campaigns to establish FZD-targeting anti-cancer drugs.

## Author contributions

Designing research studies: LG, GD, GS, PK

Conducting experiments: LG, JJSK, JW, PK

Analysing data: LG, JJSK, JW, PK

Writing the manuscript: LG, GS, PK

Review and editing of manuscript: LG, JJSK, JW, KP, GD, GS, PK

## Acknowledgements

This study was supported by Karolinska Institutet, the Swedish Cancer Society (20 0264P, CAN2017/561, CAN2018/715), the Swedish Research Council (2019-01190), Knut and Alice Wallenberg Foundation (2016.0087), Novo Nordisk Foundation (NNF20OC0063168, NNF17OC0026940, NNF19OC0056122), The Lars Hierta Memorial Foundation (FO2020-0304, FO2021-0127) and the German Research Foundation (504098926 and 331351713–SFB 1324; project A06 to G. D.).

## Supporting Figure Legends

**Fig. S1:**
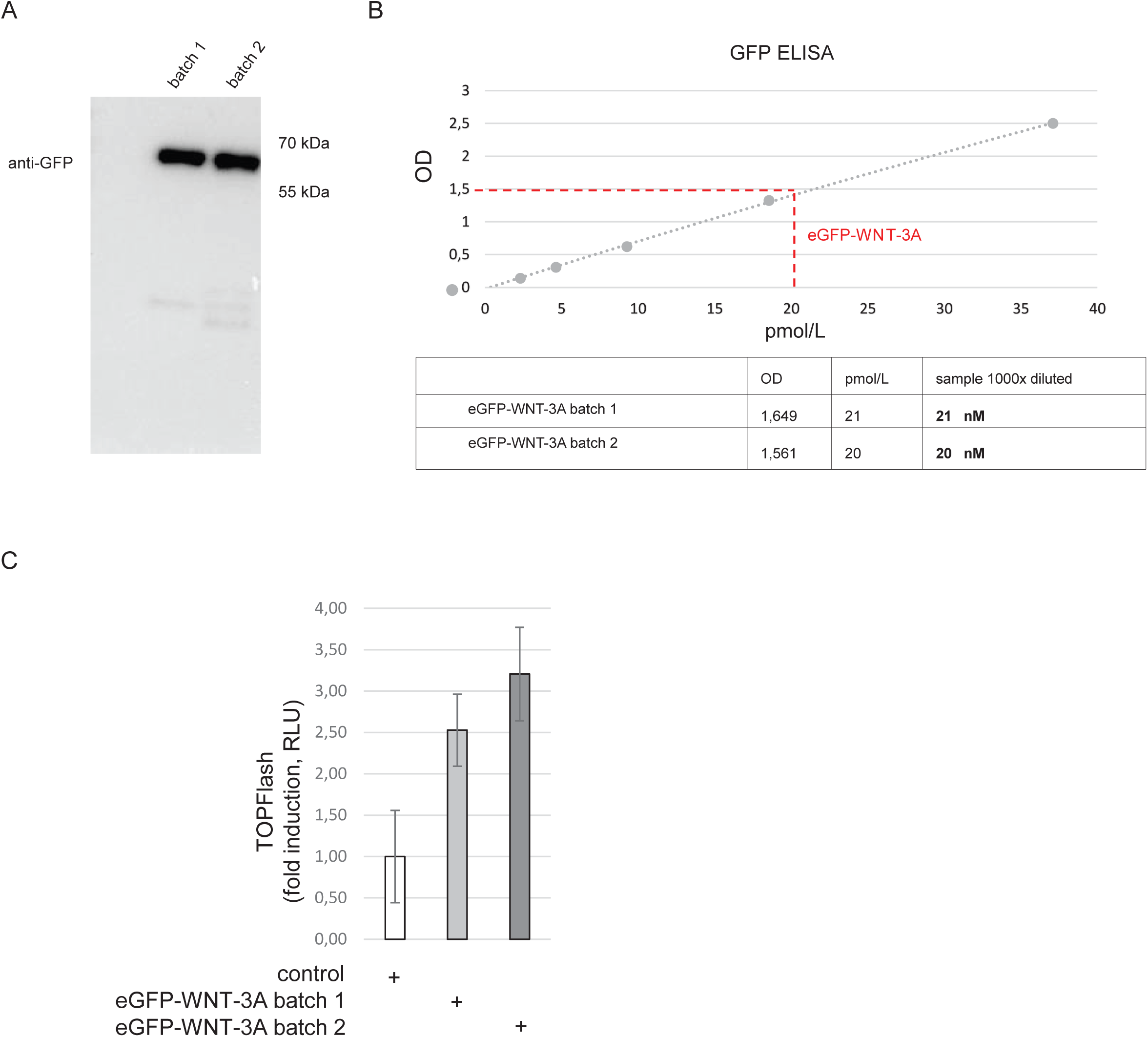
Validation of eGFP-WNT-3A conditioned media (CM). **A.** Western blotting (anti GFP, Santa Cruz Biotechnology, #sc-9996) confirms the presence of soluble full-length eGFP-WNT-3A in the CM (predicted molecular weight = 64.8 kDa). **B.** GFP ELISA assay (GFP ELISA® kit, Abcam, #ab171581) to determine the concentration of eGFP-WNT-3A present in each CM preparation. **C.** TOPFlash TCF reporter assay showing activity of the indicated CM in HEK293T cells; RLU – relative luminescence unit.

**Fig. S2:**
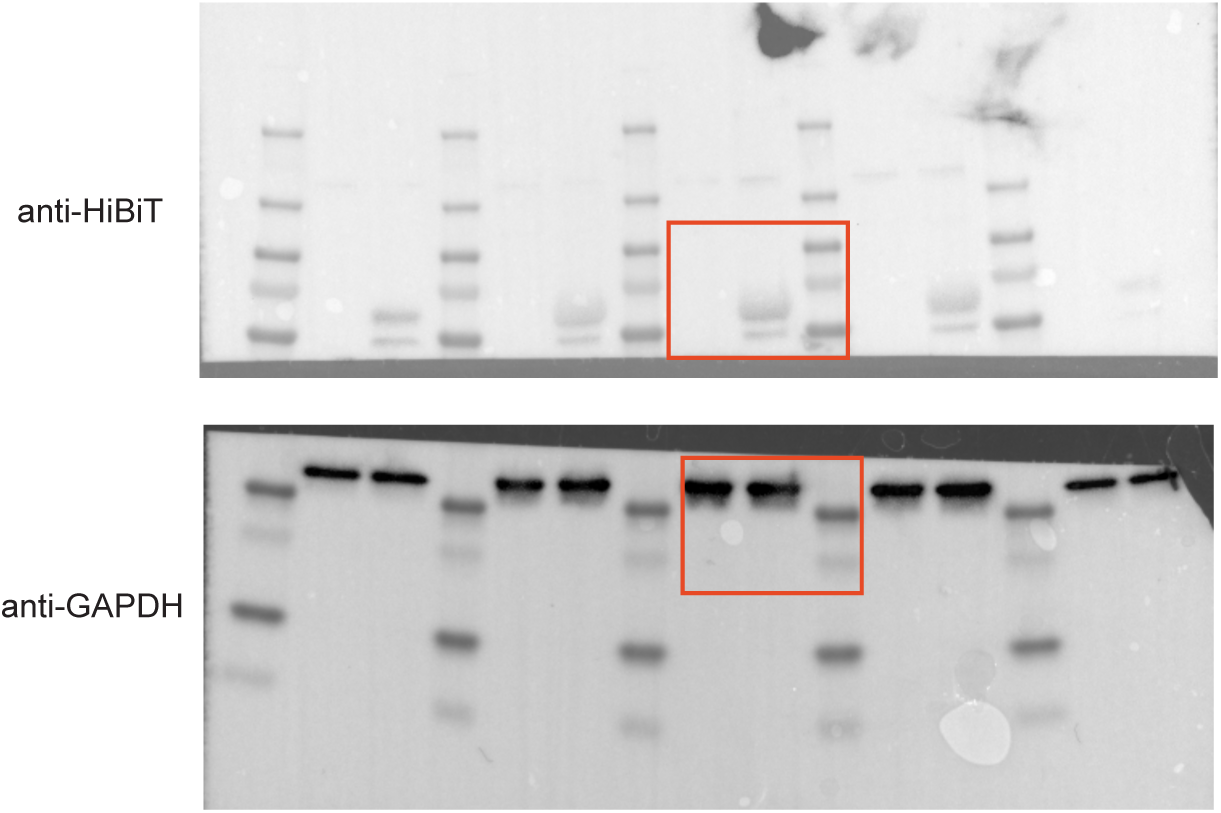
Uncropped immunoblots for HiBiT and GAPDH, which are the basis for Fig. 1. The orange boxes indicate the regions presented in the **Fig. 1**.

**Fig. S3:**
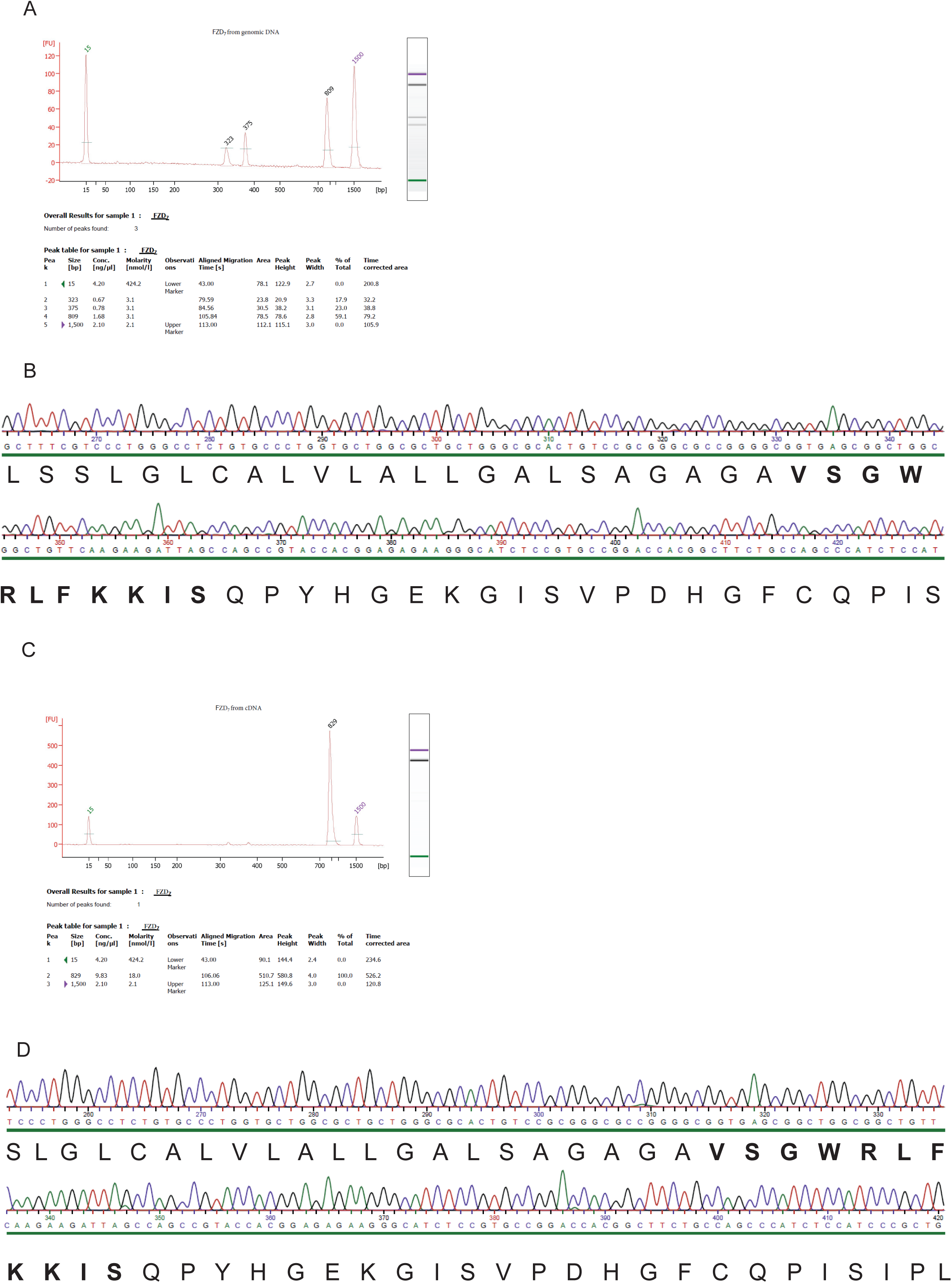
Generation of HiBiT-FZD_7_ SW480 cells with CRISPR-Cas9 genome editing. **A.** Bioanalyzer analysis of DNA products following PCR with FZD_7_-specific primers using HiBiT-FZD_7_ SW480 cells genomic DNA as a template. The predicted specifc product size = 829 bp. The sample was also subject to electrophoresis in 1% agarose gel and then the purified product (the 809 bp band) was sent for Sanger sequencing. **B.** Direct sequencing results for the PCR product from **A** show the presence one single signal. The translation from the whole part is shown under the sequencing data with HiBiT amino acid sequence marked in bold. **C.** Bioanalyzer analysis of DNA products following PCR with FZD_7_-specific primers using HiBiT-FZD_7_ SW480 cells cDNA from mRNA as a template. The predicted specific product size = 829 bp. The sample was also subject to electrophoresis in 1% agarose gel and then the purified product was sent for Sanger sequencing. **D.** Direct sequencing results for the PCR product from **C** show the presence of distinct single peaks that represent HiBiT-FZD_7_ cDNA. The translation of the whole presented sequence is shown under the sequencing data with HiBiT amino acid sequence marked in bold.

**Fig. S4:**
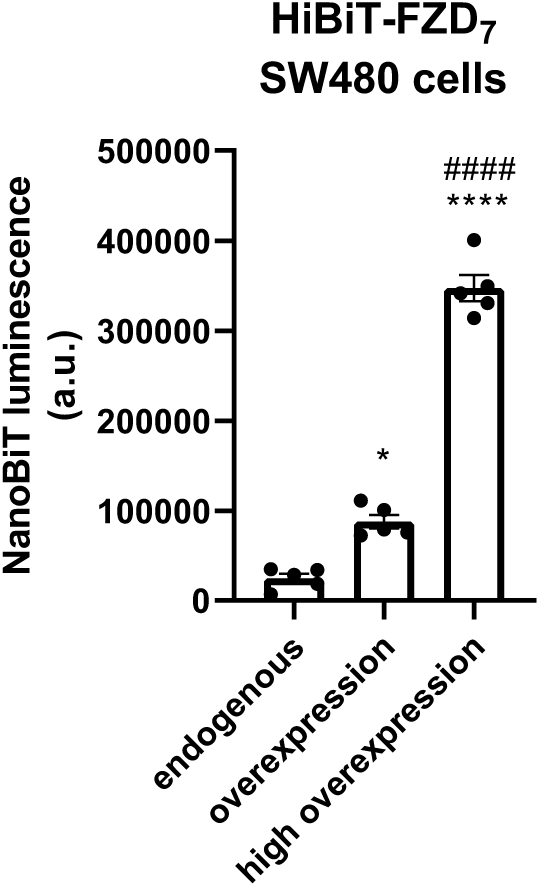
NanoBiT-emitted luminescence of the cells expressing three different FZD_7_ levels. These data were used to calculate the number of receptor molecules presented in the **Fig. 2B**. Raw data are shown from n = 5 individual experiments and are presented as mean ± SEM; a.u. -an arbitrary unit; * denotes comparison with the endogenous expression levels, # denotes comparison with the overexpressed levels.

**Fig. S5:**
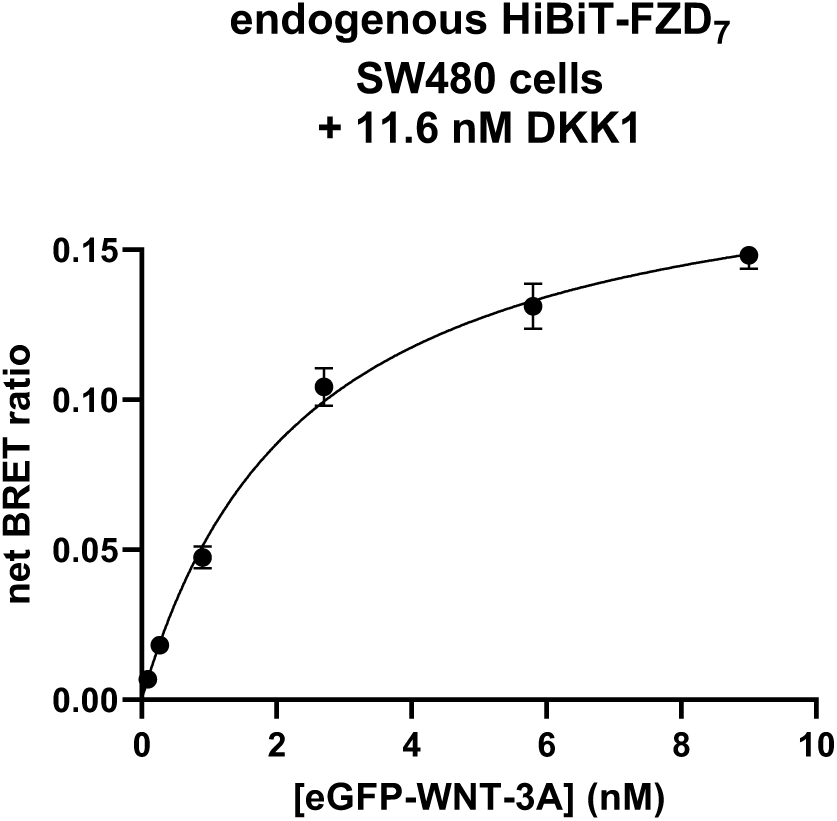
Blocking of endogenous LRP5/6 with DKK1 does not change eGFP-WNT-3A binding affinity to endogenous HiBiT-FZD_7_. Saturation binding of the eGFP-WNT-3A to endogenous HiBiT-FZD_7_ was determined using a full concentrations range (0.027 nM to 9 nM) following 180 min incubation in the presence of 11.6 nM untagged recombinant human DKK1. Raw data are presented as means ± SEM from n = 5 individual experiments, fitting a one-site specific model.

## References

1. Foord, S.M., et al., International Union of Pharmacology. XLVI. G protein-coupled receptor list. Pharmacol Rev, 2005. 57(2): p. 279–88.

2. Schulte, G., International Union of Basic and Clinical Pharmacology. LXXX. The class Frizzled receptors. Pharmacol Rev, 2010. 62(4): p. 632–67.

3. Deshpande, I., et al., Smoothened stimulation by membrane sterols drives Hedgehog pathway activity. Nature, 2019. 571(7764): p. 284–288.

4. Dijksterhuis, J.P., et al., Systematic mapping of WNT-FZD protein interactions reveals functional selectivity by distinct WNT-FZD pairs. J Biol Chem, 2015. 290(11): p. 6789–98.

5. Schulte, G. and P. Kozielewicz, Structural insight into Class F receptors - What have we learnt regarding agonist-induced activation? Basic Clin Pharmacol Toxicol, 2019.

6. Willert, K. and R. Nusse, Wnt proteins. Cold Spring Harb Perspect Biol, 2012. 4(9): p. a007864.

7. Janda, C.Y., et al., Structural basis of Wnt recognition by Frizzled. Science, 2012. 337(6090): p. 59–64.

8. Davidson, G., LRPs in WNT Signalling. Handb Exp Pharmacol, 2021. 269: p. 45–73.

9. Kozielewicz, P., A. Turku, and G. Schulte, Molecular Pharmacology of Class F Receptor Activation. Mol Pharmacol, 2020. 97(2): p. 62–71.

10. Gordon, M.D. and R. Nusse, Wnt signaling: multiple pathways, multiple receptors, and multiple transcription factors. J Biol Chem, 2006. 281(32): p. 22429–33.

11. Jung, Y.S. and J.I. Park, Wnt signaling in cancer: therapeutic targeting of Wnt signaling beyond beta-catenin and the destruction complex. Exp Mol Med, 2020. 52(2): p. 183–191.

12. Burgy, O. and M. Konigshoff, The WNT signaling pathways in wound healing and fibrosis. Matrix Biol, 2018. 68-69: p. 67–80.

13. Kozielewicz, P., et al., Quantitative Profiling of WNT-3A Binding to All Human Frizzled Paralogues in HEK293 Cells by NanoBiT/BRET Assessments. ACS Pharmacology & Translational Science, 2021. 4(3): p. 1235–1245.

14. Wesslowski, J., et al., eGFP-tagged Wnt-3a enables functional analysis of Wnt trafficking and signaling and kinetic assessment of Wnt binding to full-length Frizzled. J Biol Chem, 2020. 295(26): p. 8759–8774.

15. Schulte, G. and S.C. Wright, Frizzleds as GPCRs - More Conventional Than We Thought! Trends Pharmacol Sci, 2018. 39(9): p. 828–842.

16. Nusse, R. and H. Clevers, Wnt/beta-Catenin Signaling, Disease, and Emerging Therapeutic Modalities. Cell, 2017. 169(6): p. 985–999.

17. Bhanot, P., et al., A new member of the frizzled family from Drosophila functions as a Wingless receptor. Nature, 1996. 382(6588): p. 225–30.

18. Kowalski-Jahn, M., et al., Frizzled BRET sensors based on bioorthogonal labeling of unnatural amino acids reveal WNT-induced dynamics of the cysteine-rich domain. Sci Adv, 2021. 7(46): p. eabj7917.

19. Wright, S.C., et al., FZD5 is a Galphaq-coupled receptor that exhibits the functional hallmarks of prototypical GPCRs. Sci Signal, 2018. 11(559).

20. Kozielewicz, P., et al., Structural insight into small molecule action on Frizzleds. Nat Commun, 2020. 11(1): p. 414.

21. Schihada, H., et al., Deconvolution of WNT-induced Frizzled conformational dynamics with fluorescent biosensors. Biosensors and Bioelectronics, 2020.

22. Gibson, T.J., M. Seiler, and R.A. Veitia, The transience of transient overexpression. Nat Methods, 2013. 10(8): p. 715–21.

23. Mori, Y., et al., Development of an experimental method of systematically estimating protein expression limits in HEK293 cells. Sci Rep, 2020. 10(1): p. 4798.

24. Hsu, P.D., E.S. Lander, and F. Zhang, Development and applications of CRISPR-Cas9 for genome engineering. Cell, 2014. 157(6): p. 1262–1278.

25. Soave, M., et al., Detection of genome-edited and endogenously expressed G protein-coupled receptors. FEBS J, 2021. 288(8): p. 2585–2601.

26. Eckert, A.F., et al., Measuring ligand-cell surface receptor affinities with axial line-scanning fluorescence correlation spectroscopy. Elife, 2020. 9.

27. Goulding, J., et al., Subtype selective fluorescent ligands based on ICI 118,551 to study the human beta2-adrenoceptor in CRISPR/Cas9 genome-edited HEK293T cells at low expression levels. Pharmacol Res Perspect, 2021. 9(3): p. e00779.

28. Soave, M., et al., NanoBiT Complementation to Monitor Agonist-Induced Adenosine A1 Receptor Internalization. SLAS Discov, 2020. 25(2): p. 186–194.

29. White, C.W., et al., A nanoluciferase biosensor to investigate endogenous chemokine secretion and receptor binding. iScience, 2021. 24(1): p. 102011.

30. White, C.W., et al., NanoBRET ligand binding at a GPCR under endogenous promotion facilitated by CRISPR/Cas9 genome editing. Cell Signal, 2019. 54: p. 27–34.

31. White, C.W., et al., Using nanoBRET and CRISPR/Cas9 to monitor proximity to a genome-edited protein in real-time. Sci Rep, 2017. 7(1): p. 3187.

32. White, C.W., et al., CRISPR-Mediated Protein Tagging with Nanoluciferase to Investigate Native Chemokine Receptor Function and Conformational Changes. Cell Chem Biol, 2020. 27(5): p. 499–510 e7.

33. Kozielewicz, P., et al., A NanoBRET-Based Binding Assay for Smoothened Allows Real-time Analysis of Ligand Binding and Distinction of Two Binding Sites for BODIPY-cyclopamine. Mol Pharmacol, 2020. 97(1): p. 23–34.

34. Takada, R., et al., Assembly of protein complexes restricts diffusion of Wnt3a proteins. Commun Biol, 2018. 1: p. 165.

35. Dixon, A.S., et al., NanoLuc Complementation Reporter Optimized for Accurate Measurement of Protein Interactions in Cells. ACS Chem Biol, 2016. 11(2): p. 400–8.

36. Xu, L., et al., Cryo-EM structure of constitutively active human Frizzled 7 in complex with heterotrimeric Gs. Cell Res, 2021.

37. Do, M., et al., A FZD7-specific Antibody-Drug Conjugate Induces Ovarian Tumor Regression in Preclinical Models. Molecular Cancer Therapeutics, 2022. 21(1): p. 113–124.

38. Larasati, Y., et al., Unlocking the Wnt pathway: Therapeutic potential of selective targeting FZD7 in cancer. Drug Discov Today, 2022. 27(3): p. 777–792.

39. Flanagan, D.J., et al., Frizzled-7 Is Required for Wnt Signaling in Gastric Tumors with and Without Apc Mutations. Cancer Res, 2019. 79(5): p. 970–981.

40. Ueno, K., et al., Frizzled-7 as a potential therapeutic target in colorectal cancer. Neoplasia, 2008. 10(7): p. 697–705.

41. Vincan, E., et al., Frizzled-7 receptor ectodomain expression in a colon cancer cell line induces morphological change and attenuates tumor growth. Differentiation, 2005. 73(4): p. 142–53.

42. Tran, B.M., et al., Frizzled7 Activates beta-Catenin-Dependent and beta-Catenin-Independent Wnt Signalling Pathways During Developmental Morphogenesis: Implications for Therapeutic Targeting in Colorectal Cancer. Handb Exp Pharmacol, 2021. 269: p. 251–277.

43. Flanagan, D.J., et al., Frizzled7 functions as a Wnt receptor in intestinal epithelial Lgr5(+) stem cells. Stem Cell Reports, 2015. 4(5): p. 759–67.

44. Vincan, E., et al., Frizzled-7 dictates three-dimensional organization of colorectal cancer cell carcinoids. Oncogene, 2007. 26(16): p. 2340–52.

45. Willert, K.H., Isolation and application of bioactive Wnt proteins. Methods Mol Biol, 2008. 468: p. 17–29.

46. Kleivi, K., et al., Genome signatures of colon carcinoma cell lines. Cancer Genet Cytogenet, 2004. 155(2): p. 119–31.

47. Hulme, E.C. and M.A. Trevethick, Ligand binding assays at equilibrium: validation and interpretation. British Journal of Pharmacology, 2010. 161(6): p. 1219–1237.

48. Hoare, S.R.J., The Problems of Applying Classical Pharmacology Analysis to Modern In Vitro Drug Discovery Assays: Slow Binding Kinetics and High Target Concentration. SLAS Discov, 2021. 26(7): p. 835–850.

49. Flanagan, C.A., GPCR-radioligand binding assays. Methods Cell Biol, 2016. 132: p. 191–215.

50. Boursier, M.E., et al., The luminescent HiBiT peptide enables selective quantitation of G protein-coupled receptor ligand engagement and internalization in living cells. J Biol Chem, 2020. 295(15): p. 5124–5135.

51. Felce, J.H., et al., Receptor Quaternary Organization Explains G Protein-Coupled Receptor Family Structure. Cell Rep, 2017. 20(11): p. 2654–2665.

52. Nile, A.H., et al., A selective peptide inhibitor of Frizzled 7 receptors disrupts intestinal stem cells. Nat Chem Biol, 2018. 14(6): p. 582–590.

53. Pode-Shakked, N., et al., Resistance or sensitivity of Wilms’ tumor to anti-FZD7 antibody highlights the Wnt pathway as a possible therapeutic target. Oncogene, 2011. 30(14): p. 1664–80.

54. Fernandez, A., et al., The WNT receptor FZD7 is required for maintenance of the pluripotent state in human embryonic stem cells. Proc Natl Acad Sci U S A, 2014. 111(4): p. 1409–14.

